# Structure of a dominant negative mutant reveals a stalled intermediate state of anthrax protective antigen pore maturation

**DOI:** 10.1101/2020.06.15.152231

**Authors:** Harry Scott, Wei Huang, Srinivas Gonti, Kaiming Zhang, Nurjahan Mehzabeen, Alexander Day, Kevin P. Battaile, Scott Lovell, James G. Bann, Derek J. Taylor

## Abstract

Anthrax is a severe bacterial infection caused by *Bacillus anthracis*, which produces a tripartite toxin that includes protective antigen (PA), lethal factor (LF) and edema factor (EF). A series of dominant-negative mutations have been previously identified that prevent the heptameric PA prepore from forming the pH-induced, membrane spanning beta-barrel pore that is required for translocation of EF and LF to the cytoplasm of the infected cell. Here we show that the dominant negative D425A mutation stalls the formation of the pore at a reversible intermediate maturation state, which exhibits many of the structural aspects of the pore but fails to form the phi(ϕ)-clamp and beta-barrel structure needed for full pore maturation. Overall, this structure reveals that ϕ-clamp and beta-barrel pore formation are later steps in the pathway to pore formation, thereby providing a regulatory mechanism to prevent premature translocation of EF and LF.

## Introduction

*Bacillus anthracis*, the causative agent of anthrax, secretes an AB toxin comprised of three separate proteins: protective antigen (PA), edema factor (EF) and lethal factor (LF) (Young and Collier, 2007). Pathogenesis of the toxin requires that the 83 kDa PA component bind to host cellular receptors where it is proteolyzed to a 63 kDa form that spontaneously self-associates into a donut-shaped heptameric or octameric prepore complex (Bradley et al., 2001; Kintzer et al., 2009; Lacy et al., 2004; Scobie et al., 2003; Wigelsworth et al., 2004). Formation of the PA prepore creates high affinity docking sites for EF and LF (Melnyk et al., 2006; Mogridge et al., 2002). Binding of EF/LF facilitates endocytosis of the toxin into cells, most likely dendritic cells or macrophages of the host immune system (Abrami et al., 2003; Agrawal et al., 2003; Beauregard et al., 2000; Singh et al., 1999). Upon endocytosis, the toxins are trafficked to endosomes, and the eventual acidification of the endosomes promotes a large conformational change in PA in which the domain 2 β2-β3 loop regions of adjacent protomers form an extended β-barrel pore that spans the endosomal membrane (Abrami et al., 2004; Benson et al., 1998; Blaustein et al., 1989; Jiang et al., 2015; Katayama et al., 2010; Nassi et al., 2002; Rainey et al., 2005). In the fully mature state, F427 from each protomer extends into the lumen to form the phi(ϕ)-clamp, a ring of phenylalanine side-chains that form a narrow (6 Å) iris in the pore state (Krantz et al., 2005). The formation of the β-barrel pore and the integrity of the ϕ-clamp structure are critical for translocation of EF, a calcium/calmodulin-dependent adenylate cyclase, and LF, a zinc-metalloproteinase, into the host cell cytosol where they function to severely depress the host immune response (Bourne et al., 1974; Ghosal et al., 2017; Vitale et al., 1998).

Several residues of PA, particularly those within the D425-F427 loop, are highly conserved across *Bacillus* and *Clostridial* species and, therefore, highlight a strict requirement for a role in the structure and function of the pore (Melnyk and Collier, 2006). In 2001, Sellman and coworkers identified unique mutations that block translocation of EF and LF into the cell cytosol, including individual point mutations at three distinct residues: K397, D425 and F427 (Sellman et al., 2001). These mutations constituted dominant-negative phenotypes, where a single point mutation within each protomer of the heptamer was required to block translocation. K397A and K397D mutations prevent formation of the SDS-resistant heptamer (an assay used to show β-barrel pore formation). K397Q and K397R mutants are functional in the ability to form a pore, but are impaired in the ability to translocate an LF-fusion protein into cells. The point mutant F427A is also sufficient to block the formation of an SDS-resistant heptamer, while mutations of F427 to R, K or G strongly inhibit conversion to the SDS resistant heptamer (Sun et al., 2008). Seemingly any mutation at D425 blocks pore formation and prevents SDS resistance, whereas mutation to the adjacent residue (D426A) does not disturb the ability to form an SDS-resistant heptamer (Sellman et al., 2001).

In this study, we determined the structure of the dominant-negative D425A mutant PA oligomer at pH 5.5 using cryo-electron microscopy (cryo-EM) to a resolution of 2.7 Å. The structure identifies a unique intermediate-like state of pH-induced PA pore maturation that shares molecular features with the WT prepore and pore. The structural investigation is complemented with circular dichroism analyses (CD), kinetic light-scattering, and fluorescence analyses to reveal that the D425A intermediate is fully reversible and dependent on pH. Altogether, our investigation provides a first glimpse into the early and reversible stages of the anthrax prepore-to-pore transition and identifies the molecular deficiencies associated with the D425A mutation in pH-induced maturation events.

## Results

### Biochemical analysis of D425A heptamer identifies reversible pH-induced structural rearrangements

Each of four, distinct structural domains of a PA monomer contributes to unique functions in prepore assembly and pore maturation (Petosa et al., 1997) (Figure 1A). Despite the availability of two end-state structures for wild type (WT) PA prepore and pore (Jiang et al., 2015; Lacy et al., 2004), the extreme differences in their organization and molecular interactions has make it challenging to rationalize the sequential steps that are involved in the maturation process. Significant structural differences between the two end states include the formation of a 105 Å membrane-spanning channel, the nearly complete loss of density for domain 4 in the mature pore, and a myriad of contacts that are broken and established during the pore maturation process (Jiang et al., 2015; Lacy et al., 2004) (Figure 1B). We therefore used the D425A mutant and WT heptamers to identify differences associated with mutation that might help delineate the steps involved in the transition of the anthrax prepore to the pore state.

**Figure 1.**
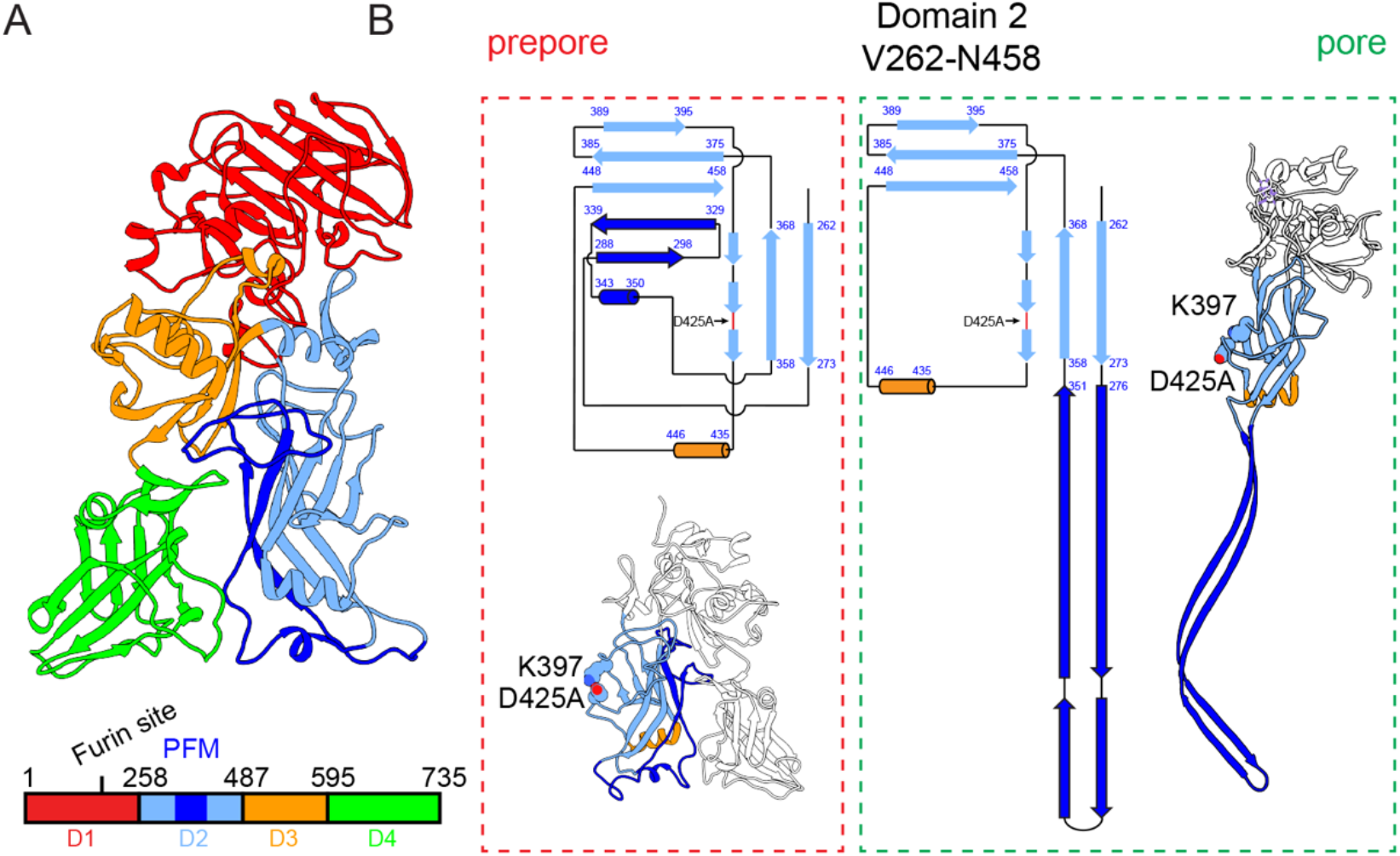
Domain organization of PA monomer. (**A**) Monomeric domain architeture and organization of PA_83_ (PDB entry: 4EE2) (Kintzer et al., 2012). (**B**) 2D schematic of the secondary structure organization of domain 2 (segment V_262_-N_458_) in the prepore (red dash box) and the pore (green dash box) states.

An SDS-resistant state associated with the mature, WT pore is not recapitulated in the context of the D425A mutant complex (Sellman et al., 2001) (Figure1-figure supplement 1). This observation indicates that the D425A mutant prevents prepore-to-pore maturation. However, initial circular dichroism (CD) studies of the heptameric prepore as a function of pH suggested that the D425A undergoes a conformational change at low pH similar to the WT heptameric protein (Figure 2). The spectroscopic changes from pH 8.5 to pH 5.5 in the CD profiles of the D425A and WT heptamers suggest a loss in alpha-helical structure (Micsonai et al., 2015). Although the CD spectra are nearly identical for both complexes at pH 8.5, those recorded at pH 5.5 are quite different for the two complexes, potentially indicating differences in secondary structure. Specifically, the WT CD spectrum has minima at 211 and 218 nm, whereas minima for the D425A heptamer are at 211 and 222 nm. Additionally, the CD minima values at pH 5.5 are significantly stronger for the WT protein (−9500 deg cm^2^ dmol^−1^ at 218 nm) than for the D425A mutant heptamer (−4400 deg cm^2^ dmol^−1^ at 222).

**Figure 2.**
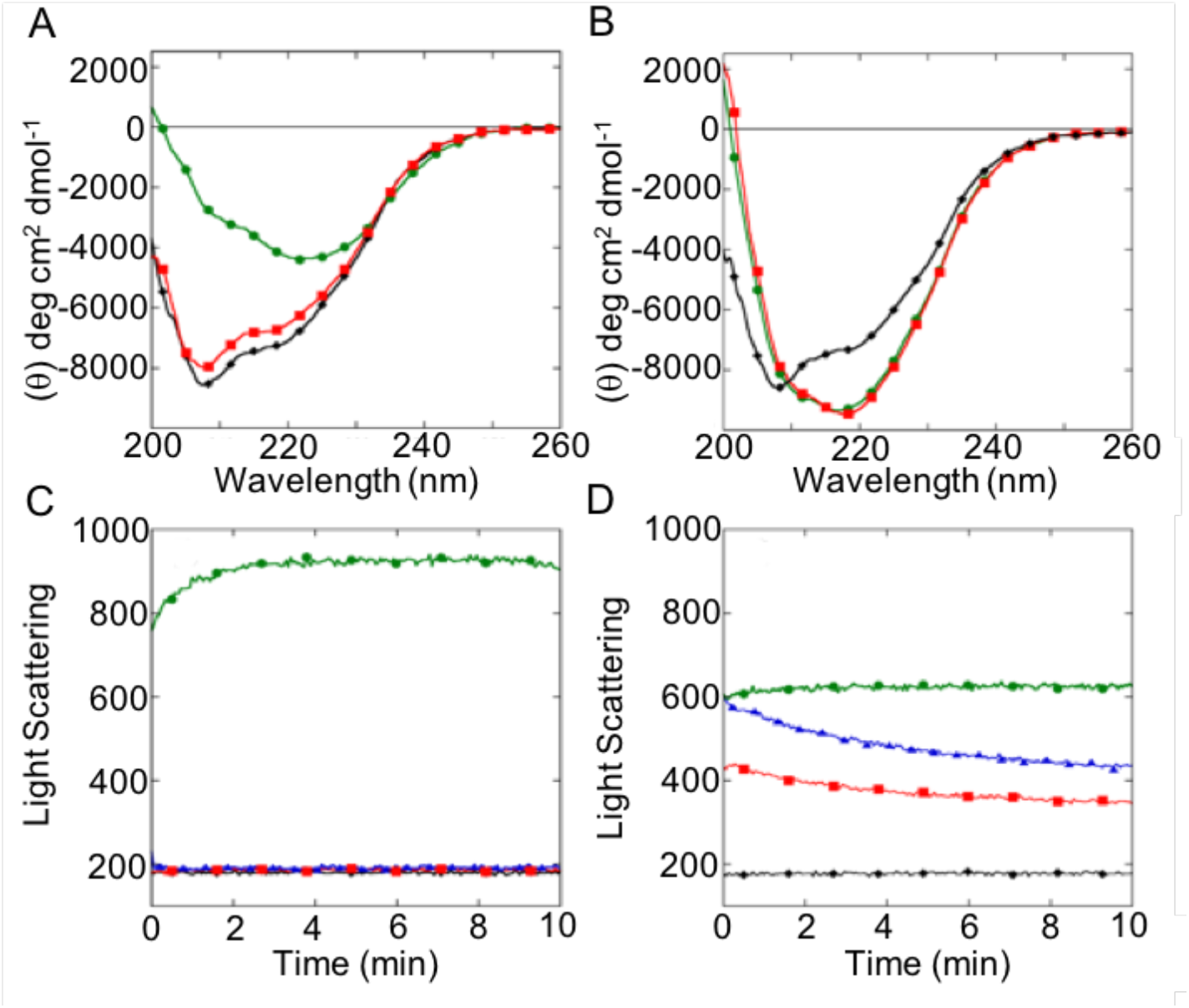
CD and light scattering experiments identify reversible pH dependent structural changes in the D425A heptamer. (**A**) CD spectra of D425A and (**B**) WT heptamer at pH 9 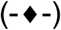, pH 5 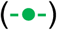 and pH 5 then back to pH 8.5 after 10 minutes 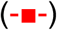. (**C**) Kinetics of light scattering measured by excitation/emission at 350 nm for D425A and (**D**) WT heptamer, under the same conditions as descried in panels **A** and **B** except at one-tenth the protein concentration. The additional blue line 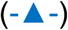 represents data where the reversion back to pH 8.5 was preceded by a 30-minute wait period at pH 5.

We next determined whether and of the pH-induced changes for WT or D425A heptamer were reversible. Reversion of the pH from 5.5 back to 8.5 resulted in a CD spectrum for the D425A heptamer that was very similar to the initial spectrum recorded at pH 8.5 (Figure 2A). This was not the case for the WT heptamer, as the spectrum recorded after a return to pH 8.5 did not change from that recorded at pH 5.5 (Figure 2B). This latter observation is consistent with the notion that the WT pore state is unable to revert back to the prepore state, as corroborated by SDS-resistance assays (Miller et al., 1999). In kinetic light scattering experiments, reversing the pH to 8.5 also reduced the degree of light scattering for both proteins (Figure 1C and D), with a near complete reversal of light scattering only for the D425A heptamer. Similar trends were observed for both complexes when the pH 5.5 state was maintained for 10 or 30 minutes before reversion back to a pH 8.5 state. Taken together, these data suggest that the initial conformational change observed upon lowering the pH to 5.5 is similar for both WT and D425A heptamers. However, the D425A heptamer stalls at a state that is fully reversible as a function of pH, whereas the WT matures to the irreversible, SDS-resistant, pore state.

### Structural characterization of the D425A heptamer

Next, we determined the three-dimensional (3D) structure of the D425A heptamer. We first attempted to crystallize D425A under different pH conditions, and were successful in growing diffracting crystals at pH 8.5. Despite the moderate-resolution (at 5.5 Å) of the resolved electron density map, it is clear that the overall structural folds of D425A mutant and the WT prepores are nearly perfectly superimposable, with a Cα root mean square deviation (RMSD) of 1.45 Å (Figure 3-figure supplement 2). This finding is consistent with studies showing that functional aspects of D425A, including LF and receptor binding, are retained (Sellman et al., 2001). Next, we determined the three-dimensional (3D) structure of the D425A intermediate at low pH (5.5) using cryo-EM and single-particle reconstruction techniques. Initially, only top-down views of the heptamer were visible due to preferred orientation on the grid; thus, a variety of conditions were tested to obtain side-views of the complex. We found that addition of 100 nM fluorinated Fos-Choline-8 helped to obtain more conformations of the complex so that the three-dimensional structure could be determined. Under these conditions, reference-free 2D classification was used to identify two distinct groups of particles that existed within the dataset. One group was indicative of a single heptameric species, while the second group represented a symmetric dimeric-heptamer structure (Figure 3-figure supplement 3).

Three-dimensional classification of these particles resulted in the identification of three distinct conformational states that included a single heptamer species resolved with C7 symmetry to 3.28 Å and two distinct dimeric-heptamers resolved with D7 symmetry to 2.67 Å and 2.80 Å, respectively (Figure 3A & B and Figure 3-figure supplement 4). The dimer interface involves the EF and LF loading platform of each heptamer, with the two structures differing in rotation of the individual heptamers that comprise the dimers. Notably, the N-terminus of each heptamer extends to occupy the substrate docking surface known as the α-clamp site (Feld et al., 2012), of the opposite heptamer (Figure 3B; Video 1). Aside from the noted differences in rotation, the RMSD for all modeled protomers (within individual or dimeric heptamers) was less than 1 Å for well-resolved regions, suggesting that the conformation of individual subunits within all three structures is remarkably similar (Figure 3C & D).

**Figure 3.**
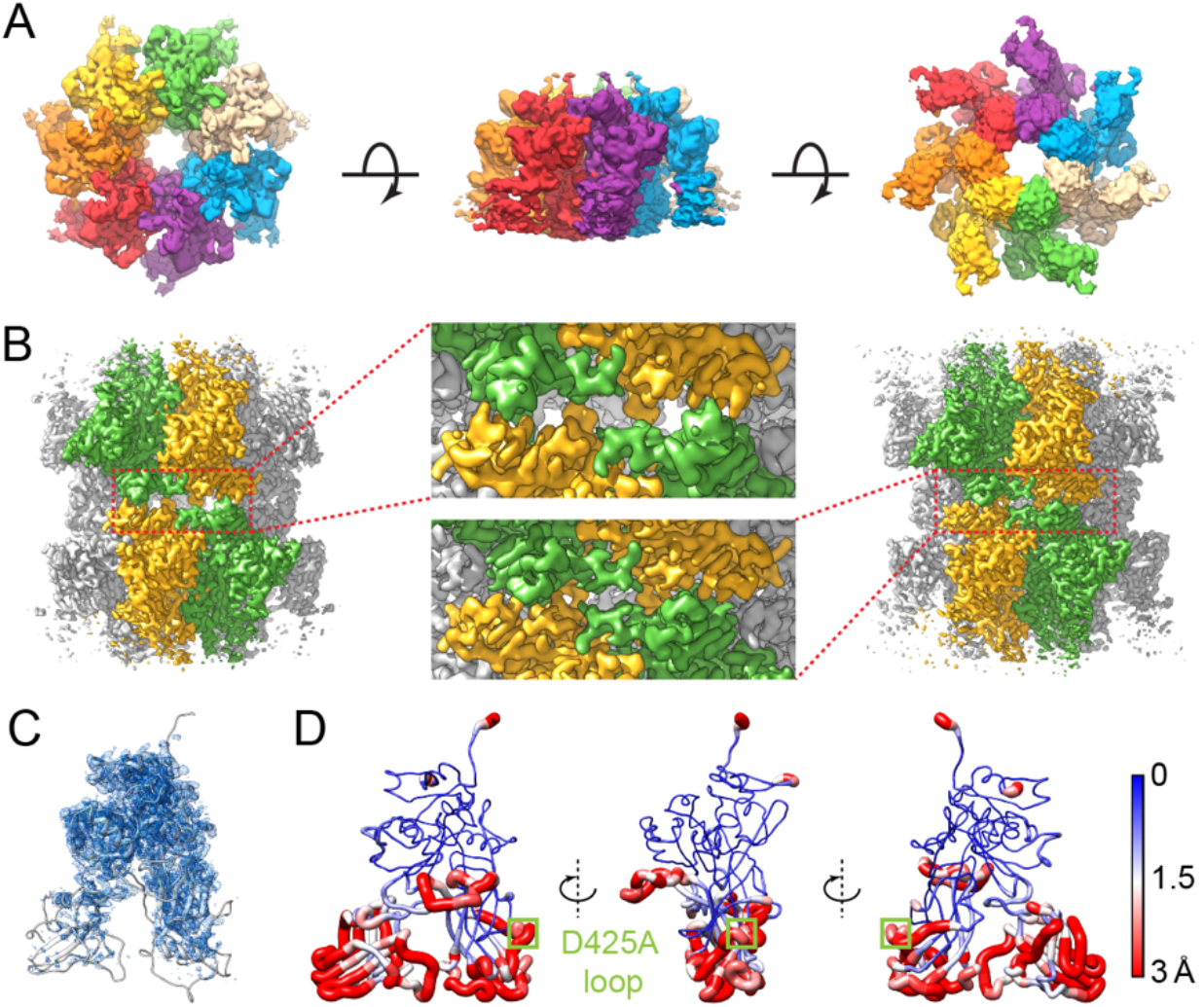
Cryo-EM analysis of the D425A intermediate structure at pH 5.5. (**A**) Overall cryo-EM 3D reconstruction of heptameric D425A intermediate at pH 5.5 shown in top, side and bottom views with individual protomers in distinct colors. (**B**) Two conformational states identified as dimers of heptameric D425A intermediates differ by interactions that form the platform for loading EF and LF. These two conformational states display distinct dimer packing interfaces highlighted in the center panels. (**C**) The cryo-EM map (blue mesh) of one protomer with a single refined structure shown in the ribbon representation in white. Domain 1 and 3 have the most well resolved map, the pore-forming module of domain 2 and majority of domain 4 correlate with weak densities with symmetry (C7/D7) applied refinement. (**D**) The sausage model of the ensemble structures (100 independently refined structures for each map) refined against three separated cryo-EM maps from one single D425A sample at pH 5.5. The root-mean-square deviation of Cα atoms is proportional to the radius of the tube and color coded as shown in the color bar.

**Video 1.** Swirling motion revealed by morphing between two heptamer dimer structures.

The cryo-EM structure of the D425A complex at pH 5.5 unveils unique features that differ from those of the wild type prepore and pore organizations (Figure 3-figure supplement 5). Specifically, the D425A structure includes detachment of domain 2 from domain 4, and the reorganization of domain 2 results in an 18% constriction of the heptamer inner lumen diameter as compared to that of the prepore (Figure 3-figure supplement 5A & Video 2). The motion of domain 2 toward the interior of the lumen coincides with a loss of intramolecular interactions with domain 4, which results in poor density for domain 4, a feature that is shared with the structure of the wild type pore (Jiang et al., 2015). Application of a low-pass filter to 12 Å of the D425A structure restored density assigned to domain 4, indicating that the loss of interactions with domain 2 allows it to shift and rotate as a folded domain, rather than being denatured at low pH (Figure 3-figure supplement 6). This notion is supported by prior studies showing that domain 4 maintains a similar fold at high and low pH (Williams et al., 2009).

**Video 2.** Conformational changes associated with PA heptamer maturation from the prepore to the D425A intermediate state. Top (left) and side (right) views display simultaneous linear morphs starting at the initial frame (prepore) and progressing to the final frame (D425A intermediate) of the MD simulation.

Maturation of the WT pore includes the formation of new stabilizing contacts between adjacent monomers that are not present in the prepore (Figure 1). For example, a key interface forms between the triple-stranded β-sheet (2β_448-458_, 2β_375-385_ and 2β_389-395_) of one protomer and the A_417_-A_420_ loop of its neighboring protomer in the mature pore (Jiang et al., 2015). This particular feature is conserved in the D425A structure, indicating that this interaction is formed in the initial stages of pore maturation (Figure 3-figure supplement 5C). Additionally, the PFM (E_275_-L_352_) of the prepore is well ordered and composed of two beta-sheets (2β_288-298_ and 2β_329-339_) with the membrane insertion loop (MIL; V_303_-F_324_) forming contacts with an adjacent protomer (Lacy et al., 2004). Maturation to the wild type pore requires the PFM to undergo a large conformational rearrangement whereby intra and inter-protomer interactions are lost, resulting in the formation of beta-barrel structure that penetrates the cell membrane (Jiang et al., 2015). The beta-barrel structure does not form in the D425A intermediate, and density corresponding to the PFM is not well resolved in the 3D reconstruction. This observation suggests that the PFM becomes disordered, and is consistent with studies that suggest that the PFM contains a pH sensor that is responsible for an initial pH-sensitive barrier that leads to the formation of an intermediate state (Kintzer et al., 2012).

### The pore maturation pathway is influenced by the conformation of individual protomers

We noticed in the single particles that in some instances there was stronger electron density for domain 4 within a heptamer compared to other protomers, suggesting that there might be incomplete conversion that was averaged out due to symmetry operations in the refinement process. To better correlate structural changes observed in PA protomers with maturation, we solved the structure of the D425A heptamer at pH 5.5 in the absence of symmetry operations. As before, both the monomer and dimeric-heptamer populations were used to reconstruct two independent volumes that were structurally very similar. The density maps identified clear differences among the protomers within each heptamer, with those on one side of the heptamer having stronger density for domain 4, while those on the opposite side correlated with weaker density for this domain (Figure 3-figure supplement 3). We next implemented a localized 3D classification approach (Ilca et al., 2015; Roh et al., 2017). First, two density masks were generated; one for the whole refined heptamer map and the other for a single protomer within the assembled heptamer. The two maps were applied to the heptamer and the difference yielded a soft mask that was applied to subtract density outside of the individual monomer. This mask was applied to the refined heptamer map to generate the density map for calculating the projected images to be subtracted from the raw particle images. The operations were extended to include all seven protomers within the assembled heptamer, which resulted in an expanded dataset, with seven or 14 times the number of individual units, with sub-particle images maintaining signals from a single protomer (Figure 4A). This analysis resulted in a nearly equal distribution of seven distinct protomer structures that could be identified within the assembled heptamer.

**Figure 4.**
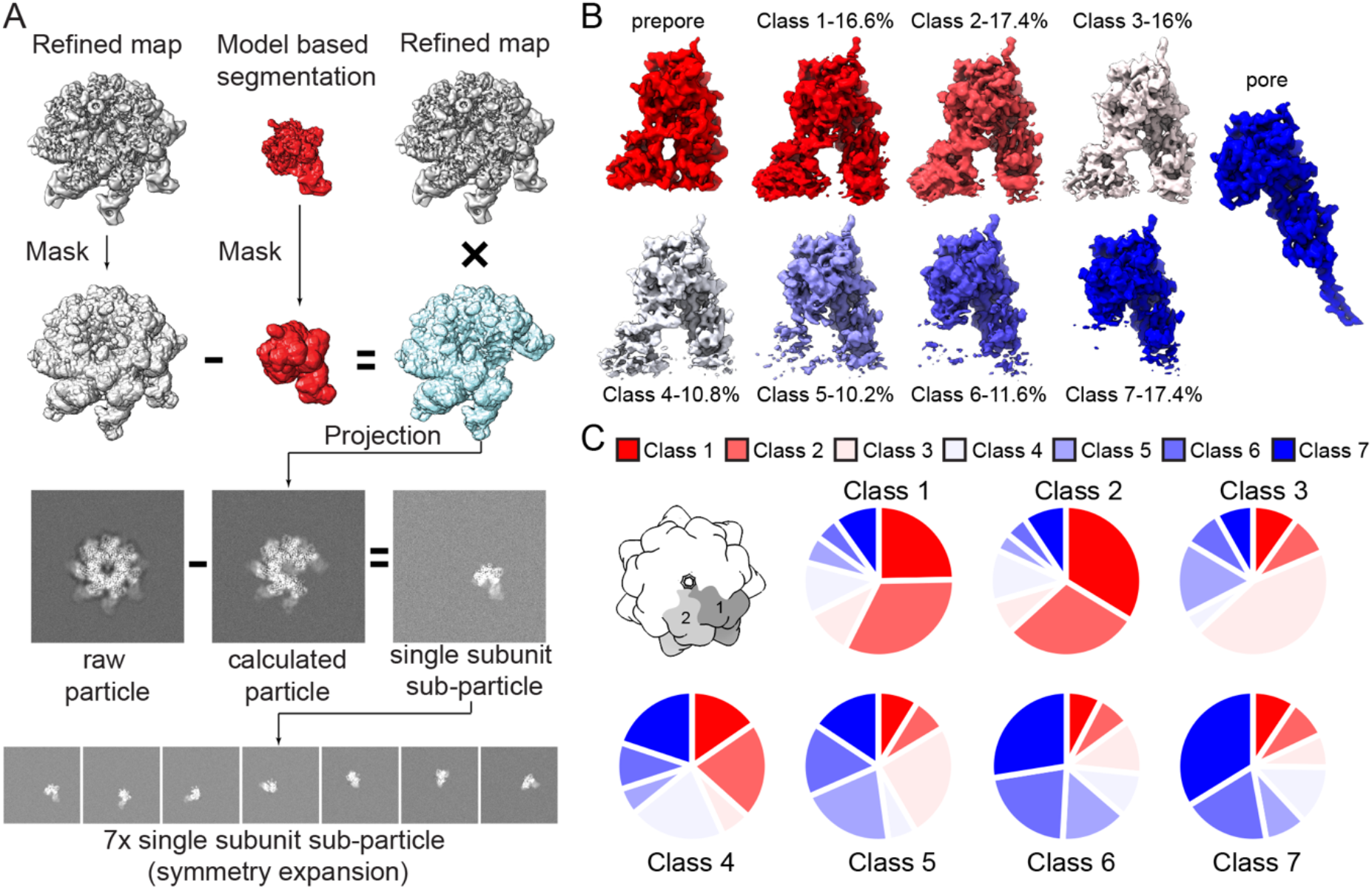
Focus classification of individual protomers identifies structural heterogeneity. (**A**) Schematic of the image analysis workflow for generating the raw images for single protomer sub-particles using symmetry expansion and image subtraction based on projected experimental density. (**B**) Cryo-EM densities of the individual classes along with the prepore and pore maps, shown with the core scaffold at a similar density level. The population of each class is shown at the top. The prepore map is calculated using PDB entry 1TZO (Lacy et al., 2004), and the pore map EMD-6224 (Jiang et al., 2015). (**C**) Probabilities of particles in each class next to the other classes shown as pie charts. For example, when position 1 resides as a particle from class 1, the particle at position 2 is more likely to belong to class 1 and class 2.

The seven protomer structures were defined as class 1 having the strongest density for domain 4 and, therefore, being most similar to the wild type prepore structure, and class 7 having the weakest density for domain 4 and most closely resembling the pore state (Figure 4B). Classes 2-6 were defined by decreasing density for domain 4 (Video 3). The number of distinct protomer structures within each D425A heptamer varied among the individual complexes. Therefore, the influence of a precise conformation for an individual protomer was compared to that of the neighboring protomers. We calculated the distribution of classes that each single protomer was assigned in the local focused classification step for the seven pairs of neighboring protomers within a heptamer. This analysis identified a proximity based structural influence for protomers within the heptamer as the conformation of individual protomers was structurally most similar to that of adjacent protomers (Figure 4C). This result indicates that the pH-induced maturation of individual protomers is influenced by the state of adjacent protomers, and suggests that complete maturation proceeds in a directional manner analogous to a domino toppling effect.

**Video 3.** Morphing of the density maps for individual monomers analyzed from the asymmetric reconstruction illustrates the trajectory of transition starting from the prepore state (red) then advancing through seven distinct classes from the localized classification of our intermediate state and ending at the WT pore state (blue).

All seven protomer structures displayed a shift of domain 2 toward the interior of the assembled heptamer that is similar in scale to that of the symmetrically averaged reconstruction and of the WT pore structure. The primary structural differences among monomers within the assembled asymmetric heptamer can be attributed to the gradual, but coincident, loss of density for domain 4 and the PFM in domain 2, even though the contacts between domain 2 and domain 4 have already been lost in each individual protomer (Video 3). Those classes with stronger domain 4 density were associated with stronger density for the lower part of domain 2 attributed to a region of the PFM (the 343-350 loop), which was missing in the symmetric reconstruction (Figure 4-figure supplement 7). This region forms a helix (2α_343-350_) in the prepore state that undergoes a secondary structure transition during pore maturation to form part of the membrane-spanning β-barrel in the pore state ^18^ (Figure 1B). At low pH this helix becomes disordered before the multi-domain β-barrel forms in the mature, wild type pore (Chadegani et al., 2014; Petosa et al., 1997; Rajapaksha et al., 2012). As this particular helix also participates in receptor binding of the prepore (Jiang et al., 2015; Lacy et al., 2004; Scobie et al., 2007), the structural rearrangement likely contributes to the dissociation of the pore from the receptor. A comparison of the corresponding density for this region in the seven individual monomeric structures suggests that it moves toward the interior of the pore during the initial stages of maturation (Figure 4-figure supplement 7). That the electron density for this region is better resolved in the classes that have stronger density for domain 4 strongly indicates that the pH-induced shift of domain 2 away from domain 4 is coincident with destabilization of this helix. Interestingly, a monoclonal antibody binds to this precise helix to prevent pore formation, thereby further highlighting a critical role of this helix in the pore-maturation cascade (Mechaly et al., 2012).

## Discussion

The toxicity of the anthrax tripartite system relies almost entirely on the ability of the prepore to form the pore so that EF and LF can be translocated to the cytoplasm of the infected host (Janowiak et al., 2009). The wild type PA prepore state is inactive, with an inability to translocate these toxins, and the PFM is stabilized by intra- and intermolecular interactions. Our CD analysis shows that the pH 8.5 structures of the prepore are similar, and that at pH 5.5 the structures are quite similar despite differences in minima. For instance, it may be that in the WT structure, the CD minimum of 218 could suggest a greater degree of beta sheet structure for the WT protein compared to D425A. Importantly, the spectroscopic data reveal that the D425A heptamer undergoes initial conformational changes that are fully reversible and pH dependent. Using cryo-EM, we show that the D425A mutant at pH 5.5 forms a structure that is different than either the prepore or pore states, and we characterize this an intermediate that is stalled along the pathway to pore formation. A shift of domain 2 toward the inner lumen, as seen in the stalled intermediate state structure, generates a new inactive conformation and represents an early transitional state, occurring before formation and stabilization of the ϕ-clamp which is a prerequisite for β-barrel formation (Jiang et al., 2015). Our data support a mechanism of anthrax PA maturation that is initiated by a pH-sensitive trigger that results in a constriction of the lumen of the heptamer that is coincident with separation of domains 2 and 4 of individual protomers. This observation is consistent with studies showing that separation of domain 2 and domain 4 is an early step in pore formation, and likely a consequence of protonation of a pH sensor that resides in the PFM (Kintzer et al., 2012). Release of the PFM of domain 2 causes this region to become more dynamic and less ordered for individual protomers within the heptameric complex. The D425A intermediate state, while undergoing much of the structural changes needed to achieve the pore, resides in a reversible intermediate state of maturation.

Our structure identifies regions of the protein that undergo pH-induced conformational changes, including the PFM and particularly the 343-350 loop from domain 2 that interacts with the host cellular receptor. In the pore structure solved by cryo-EM (Jiang et al., 2015), this region forms part of the long β-barrel stem, which presumably cannot form in the case of D425A. Although aggregation phenomena in general is poorly understood, it may be that the 343-350 region contributes significantly to aggregation at pH 5.5 where it is dynamically flexible and hydrophobic residues are exposed, and less so when the full stem structure is created in the pore.

Histidine protonation has been postulated to be involved in pore formation, but experiments have been largely inconclusive. For instance, single point mutations of histidines within the PFM do not alter the pH-sensitive barrier to pore formation, and uniform labeling of PA with 2-fluorohistidine (2-FHis), which has a side-chain pKa of ~1, does not affect pore maturation (Wimalasena et al., 2007). However, in the latter case, addition of the soluble von-Willebrand factor A domain of the receptor CMG2, which binds at the domain 2-domain 4 interface in a 2-FHis background does in fact largely prevent pore formation (Wimalasena et al., 2010). Other studies have determined that the initial pH-sensitive barrier that must be overcome in the transition to the pore is associated with separation of domains 2 and 4, a barrier that is dependent on E302(Kintzer et al., 2012). Accordingly, removal of the capability of protonation of E302 (E302A), or potentially *en masse* histidines by 2-FHis labeling coupled with receptor binding, raises the initial pH barrier required for separation of domains 2 and 4 to such an extent that pore formation is largely blocked. As such, receptor dissociation would allow domains 2 and 4 to separate, allowing the ϕ-clamp to form and pore formation to ensue.

How do mutations at D425 prevent complete pore maturation? Although other mutations have been found to be dominant negative, the effect of D425A was made strikingly clear in studies by Janowiak and coworkers, who purified (D425A_1_/WT_6_) heptamers and showed that these were unable to form pores and were non-functional (Janowiak et al., 2009). This question is particularly intriguing as the mutation of a single residue at this site (including conserved mutations Asn and Glu) entirely prevents formation of the pore. In our intermediate structure presented here and in the prepore structure, electron density in the D425 containing loop is poor, suggesting that this region is highly flexible and disordered. However, density to accommodate D425 is obvious in the wild type pore state and is located in a confined, nonionic pocket that includes intra and inter-molecular contacts with S428 and T430 (Jiang et al., 2015). Interestingly, the conservative mutation of S428 to cysteine leads to an ~100-fold reduction in cell-mediated toxicity (Mourez et al., 2003), suggesting that these intra- and inter-molecular contacts are critical for positioning of the ϕ-clamp, and that in a deprotonated state these contacts may not form or are prohibited from forming. As such, it may be that only an Asp at this residue, given its size, can accommodate this confined pocket.

Alternatively, it may be that mutations at D425 alter steps leading up to the formation of the ϕ-clamp. While mutations at K397 to Ala or Asp similarly block pore formation, mutation to Gln could be made functional by making a corresponding mutation at D426 to Gln. Melnyk and coworkers showed that while K397 and D426 were conserved as Lys and Asp in *Bacillus* species, they are conserved as Gln and Gln in *Clostridium* species. Since Gln is uncharged, it suggests that electrostatic interactions at these two sites are not a necessary requisite for pore formation. Nonetheless, it was posited that the K397 and D426 containing loops establish a structural scaffold for proper positioning of the ϕ-clamp, and we cannot rule out that any mutation at D425 also disrupts this structural scaffold needed for correct ϕ-clamp positioning. It is worth noting that the data set used for the cryo-EM structure of the mature, wild type pore included a subset of particles that was used to determine a distinct pore-like structure. This structure was characterized as having clear density for the ϕ-clamp, but with absolutely no density for the PFM (Jiang et al., 2015). These data, taken together with the structures in the intermediate state presented here, delineate the potential trajectory from the prepore state to the pore state (Figure 5).

**Figure 5.**
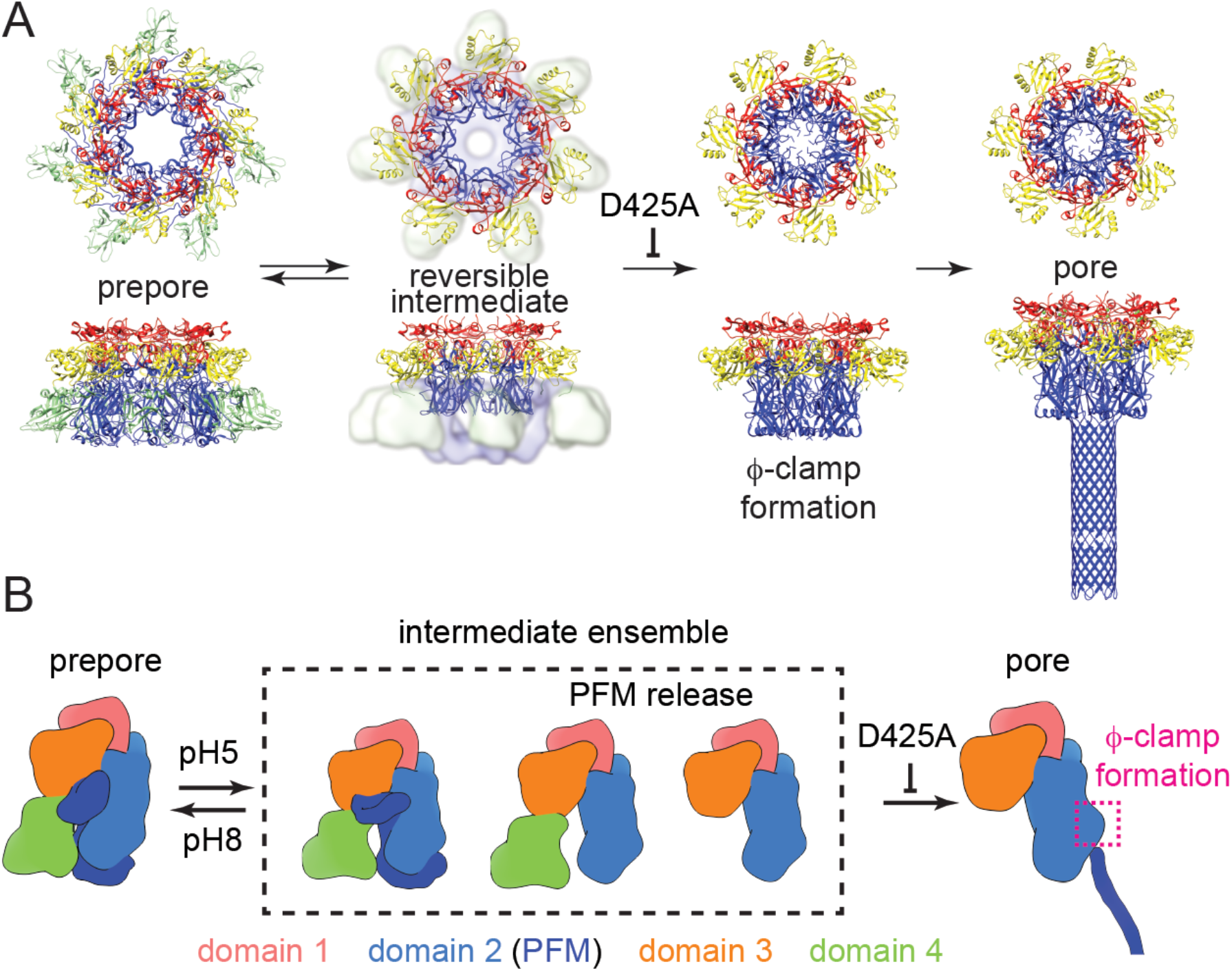
Proposed sequential events towards pore formation in the anthrax PA system. (**A**) Related structures illustrating steps towards pore formation. Prepore structure (PDB entry: 1TZO) (Lacy et al., 2004), reversible intermediate state (current study) with partial domains in molten globule state shown in low resolution density, quasi-pore (EMD-6225) (Jiang et al., 2015) and pore structures (EMD-6224 and PDB entry: 3J9C) (Jiang et al., 2015). (**B**) Stepwise mechanism showing pore formation.

Early formation of the ϕ-clamp, before the formation of the membrane penetrating pore state, has important functional consequences. The ϕ-clamp forms a leak-free seal around the translocating polypeptide, and this structure is critical for maintenance of a proton gradient from the acidic early endosome to the more neutral cell cytosol, which is needed to drive translocation (Krantz et al., 2006). This structure supports a more generalized mechanism of pore formation, whereby formation and folding of a selectivity filter occurs prior to membrane penetration, which would help prevent the dissipation of the charge gradient across the membrane (Bann, 2012; Delaney et al., 2014; Gajewski et al., 2011).

## Materials and Methods

### Protein Purification

Expression of PA and the D425A mutant PA were performed with *E. coli* UTH780 containing the plasmid pQE80-PA as described previously (Wimalasena et al., 2007). Mutation of D425A and E712C was carried out using the Stratagene Quikchange mutagenesis kit, using pQE80 PA as template, and the resulting mutant PA plasmids verified by sequencing. PA and D425A proteins were purified as described (Wimalasena et al., 2007). Purification and labeling of PA-E712C, and D425A-E712C with AF350 (Invitrogen) was carried out as described previously (Evans, 2012). To form the prepore, monomeric PA (10 mg) in 20 mM Tris pH 8, 150 mM NaCl was digested with 10 μg Trypzean (Sigma) for 30 min at RT and quenched with 100 μg soybean trypsin inhibitor. The prepore complex was isolated using a Superdex S200 size-exclusion column equilibrated in 20 mM Tris, 0.4 M NaCl, pH 8.5, 1 mM CaCl_2_.

### SDS-Resistant Pore-Formation Assay

To start the reactions, 6 μL 33.5 μM WT and 6.25 μL 32 μM D425A prepore was added to 194 μL or 193.75 μL of 50 mM Tris, 25 mM MES, 25 mM Acetate buffer at pH 5.5. 21 μL samples were taken at 2.5, 5, and 30 min and 1, 6 and 24 hr and added to 7 μL 4X SDS loading buffer to stop the reaction and heated to 70°C for 5 min. Samples were separated by gel electrophoresis using Invitrogen Tris-Acetate gradient gels. Protein bands were visualized using a Coomassie Brilliant Blue staining procedure. Analysis for the gels was performed in ImageQuant where the signal for the top two bands, representing the SDS-resistant pore species, was combined and represented as a single value and a ratio was calculated with the signal for the PA_63_ species.

### Crystallization and Data Collection

A sample of the prepore (PA_63_)_7_ (D425A) at a concentration of 15 mg/mL in 20 mM Tris pH 8.5, 0.4 M NaCl, 1mM CaCl_2_ was used for crystallization screening in Compact 300 (Rigaku Reagents) sitting drop vapor diffusion plates at 20°C using equal volumes of protein and crystallization solution equilibrated against 75 μL of the latter. Samples obtained at higher pH (10% (v/v) ethanol 100 mM Tris pH 8.5) were transferred to a fresh drop composed of 80% crystallization solution and 20% (v/v) ethylene glycol and stored in liquid nitrogen prior to data collection. These crystals yielded low resolution diffraction to 5.5 Å resolution (Supplemental Table 1). X-ray diffraction data were collected at the Advanced Photon Source beamline 17-D using a Dectris Pilatus 6M pixel array detector and the intensities were integrated using with XDS (Kabsch, 2010; Kabsch and IUCr, 1988) via Autoproc (Vonrhein et al., 2011) and the Laue class analysis and data scaling were performed with Aimless (Evans, 2011). Structure solution was conducted by molecular replacement with Phaser (McCoy et al., 2007) using a previously determined PA-prepore structure (PDB: 1TZO (Lacy et al., 2004)) as the search model. The top solution (TFZ=44.0, LLG=10529) was obtained in the space group *P*2_1_ and contained two heptamers in the asymmetric unit. The *B*-factors were set to 257 Å^2^ (Wilson *B*-factor for the data) and the model was refined with Buster (Blanc et al., 2004; Bricogne G., Blanc E., Brandl M., Flensburg C., Keller P. and Roversi P, Sharff A., Smart O.S., Vonrhein C., 2019) using non-crystallographic symmetry restraints (Smart et al., 2012). Iterative rounds of refinement and manual building were conducted with Buster and Coot (Emsley et al., 2010), respectively.

### Cryo-Electron Microscopy

The D425A prepore complex (8.43 μM – pH 8.25) was diluted 1:3 with 50 mM Tris, 25 mM MES, 25 mM Acetate and 200 mM NaCl buffer (pH 5.0) and 1 μl 900 uM F-FOS-Choline 8 (in the same buffer) was added to the sample. Holey carbon grids (Quantifoil Cu 300 mesh R1.2/1.3 grids were loaded with 3.5 μL sample and plunge frozen in liquid nitrogen cooled liquid ethane using a vitrobot (FEI) with a blotforce of −10 for 3.5 seconds at 100% humidity and 4°C. The frozen specimens were imaged at 300 keV on a Titan Krios TEM equipped with a K2 direct electron detector at a nominal magnification of 165,000 with a pixel size of 0.85 Å in counting mode. 3564 movie stacks were collected over the course of 3 days using EPU software (ThermoFisher) with a defocus range of −0.5 to −2.5 μm.

### Image Processing

Raw movies were motion corrected and dose-weighted using MotionCor2 1.0.2 (Zheng et al., 2017), the first two frames and the last frame were ignored during moving processing. Particles were automatically picked using Gautomatch (https://www.mrc-lmb.cam.ac.uk/kzhang/Gautomatch/) with a gaussian blob with a 200Å radius. Contrast transfer function parameters were estimated using Gctf (Zhang, 2016). We then performed 2D reference-free averaging using Relion 2.0 (Kimanius et al., 2016) to eliminate junk particles (Fernandez-Leiro and Scheres, 2017). The selected particles were imported into cryoSPARC (Punjani et al., 2017) to generate the initial model with D7 symmetry imposed. The initial model was used as the reference in the 3D classification carried out in Relion 2.0 for global angular search and the particles were divided into five classes with a regularization parameter T=4 and the imposition of D7 symmetry. The class with the highest resolution and the most particles was used for refinement using 55,912 particles in cisTEM (Grant et al., 2018) and subsequent map sharpening generating the final model at a resolution of 2.67 Å as defined by gold-standard fourier shell correlation 0.143 criterion. Local resolution values were calculated using RESMAP (Kucukelbir et al., n.d.).

### Model Building and Refinement

The prepore crystal structure (1TZO) with the missing regions modeled using MODELLER and an alanine residue at the D425 position (Eswar et al., 2007; Fiser et al., 2000) was docked into the cryo-EM map. Molecular dynamic flexible fitting (MDFF) was then performed using the prepore crystal structure as the starting structure using NAMD (see details in Molecular dynamic flexible fitting section) (Trabuco et al., 2008). The final frame of the MDFF trajectory was subjected to real-space refinement using PHENIX (Adams et al., 2010) with the cryo-EM map as a constraint. Regions that did not converge into the density were manually adjusted using Coot (Emsley et al., 2010). In the final model, residues with clear side-chain densities were presented with all-atom model, while the rest with only Cα model. The final all-atom model was evaluated using MolProbity (Chen et al., 2010). The statistics for these processes are described in Supplemental Table 2.

### Focused classification of localized reconstruction

First, each of the 14 subunits was aligned to the same orientation using the “relion_particle_symmetry_expand” command based on the D7 symmetry and the refined orientation parameters for the particles used to calculated the 2.67 Å map. Second, two masks for the D425A complex and a single subunit were generated using Relion with a 5 pixel radius spherical mask to dilate the mask and a 0 cosine edge padding. A mask for 13 subunits was calculated by subtracting the complex mask with the single subunit mask, then multiplied to the 2.67 Å D425A complex map to yield a new map for 13 subunits. Third, the new particle images for a single subunit were computed by subtracting the aligned particle images from the first step with projected images of the new 13-subunit map with corresponding refined orientation parameter. A new map was then calculated using these new particle images for a single subunit with “relion_reconstruct” to validate the above steps. This new set of 14 × 55,912 subparticle images were used from 3D focused classification (k = 7) without image alignment with a regularized parameter T = 16.

### Molecular Dynamics – Flexible Fitting

To determine the molecular motions required to form the pore-intermediate, the map was first converted to situs format using the Situs map2map (Wriggers, 2012) command and the prepore crystal structure (1TZO) with the missing regions modelled using MODELLER was fit into the map as a rigid body using UCSF Chimera (Pettersen et al., 2004). The situs map was subsequently converted to an MDFF potential in VMD. Using VMD, the prepore crystal structure was used to make a .psf file containing all of the connectivity and partial charge information for the structure. Secondary structure restraints were generated base on the .psf file information. MDFF was performed using NAMD (Trabuco et al., 2008) with an initial gscale of 0.3 for the first 50,000 steps and a gscale value of 10 for the minimization step.

### Circular Dichroism

CD experiments were initiated with 22 μl of 55 μM PA_63_ or D425A PA_63_ in 20 mM Tris, 400 mM NaCl, 1 mM CaCl_2_ pH 8.5. For experiments measured at alkaline pH, 28 μL of 0.2 M CHES pH 9 was initially added, followed by an additional 150 μL of 0.15 M CHES pH 9 before CD spectra were recorded at pH 9. Similarly, for experiments measured at acidic pH, 28 μL of 0.2 M MES pH 4.8 was initially added, followed by an additional 150 μL of 0.15 M MES pH 4.8 before CD spectra were recorded at a final pH of 5. For the revered pH experiments, PA at pH 8.5 was first adjusted to pH 5.5 by the addition of 28 uL of 0.2 M MES pH 4.8. After 10 min, the pH was returned to 8.5 with the addition of 150 μL of 0.15 M CHES pH 9 buffer prior to CD measurements. All CD data were recorded on a Jasco J810 spectropolarimeter using a water jacketed cylindrical cell of 0.5 mm pathlength, and recorded from 260 to 199 nm with a scan speed of 20 nm/min and a time constant of 2 s. All data represent an average of 5 scans. Buffer baselines were subtracted from the data, and data were normalized to mean residue ellipticity. The temperature was maintained at 20°C throughout the experiments. Each experiment was performed in triplicate with the data shown representing each replicate that was performed under identical conditions.

### Kinetics Light Scattering experiments

Light scattering kinetics were performed using conditions similar to those described for the CD experiments with the exception PA concentrations, which were 10-times less than those used in the CD experiments. Briefly, 2.2 μL of PA (55 μM) were mixed with 19.8 μL of 20 mM Tris, 400 mM NaCl, 1 mM CaCl_2_ to yield a final volume of 22 μL. For kinetic measurements, 22 μL of 5.5 μM PA was quickly diluted with 28 μL of 0.2M MES 4.8 or CHES pH 9, then followed by the addition of 150 μL of 0.15 M MES pH 4.8 or CHES pH 9. Reactions were mixed in a micro quartz cuvette where data acquisition were measured. For the reversible pH experiments, pH was first adjusted to pH 5 by adding 0.2 M MES pH 4.8. After 10 or 30 minutes at pH5, the reaction was returned to pH 8.5 with the addition of 0.15 M CHES pH 9. Light scattering experiments were performed by monitoring the Ex 350 nm/Em 350 nm with a Cary Eclipse spectrophotometer. The excitation/emission slits were set to 5 nm, and data were recorded with an averaging time of 0.1 s for a total of 10 min at 20°C. Each experiment was performed at least three times and the data are representative of those replicates, each replicate being an independent experiment performed under identical conditions.

## Accession Numbers

The cryo-EM reconstructions of D425A intermediate at pH 5.5 have been deposited into the Electron Microscopy Data Bank under accession codes EMD-20793, 20803, and 20804. The X-ray crystallography data of D425 at pH 8.5 have been deposited into the Protein Data Bank (PDB) under accession code 6UJI.

## Acknowledgements

We thank Dr. Wah Chiu and the Cryo-EM data collection facility consortium at National Center for Macromolecular Imaging (NCMI) and Stanford and the Stanford Linear Accelerator Center (SLAC) for access to the Titan Krios and for assistance with data collection. This study was supported by NIH grants DP2 CA186571, R01 CA240993, and R01 GM133841 (D.J.T.); a Regional Cryo-EM Data Collection Consortium Grant (U24 GM116787) at the NCMI (Chiu, MPI); and American Heart Association grant 17POST33650070 (W.H.). Use of the University of Kansas Protein Structure Laboratory was supported by a grant from the National Institute of General Medical Sciences (P30 GM110761) at the National Institutes of Health. Use of the IMCA-CAT beamline 17-ID at the Advanced Photon Source was supported by the companies of the Industrial Macromolecular Crystallography Association through a contract with Hauptman-Woodward Medical Research Institute. Use of the Advanced Photon Source was supported by the U.S. Department of Energy, Office of Science, Office of Basic Energy Sciences, under Contract No. DE-AC02-06CH11357.

---

**Figure 1-figure supplement 1.**
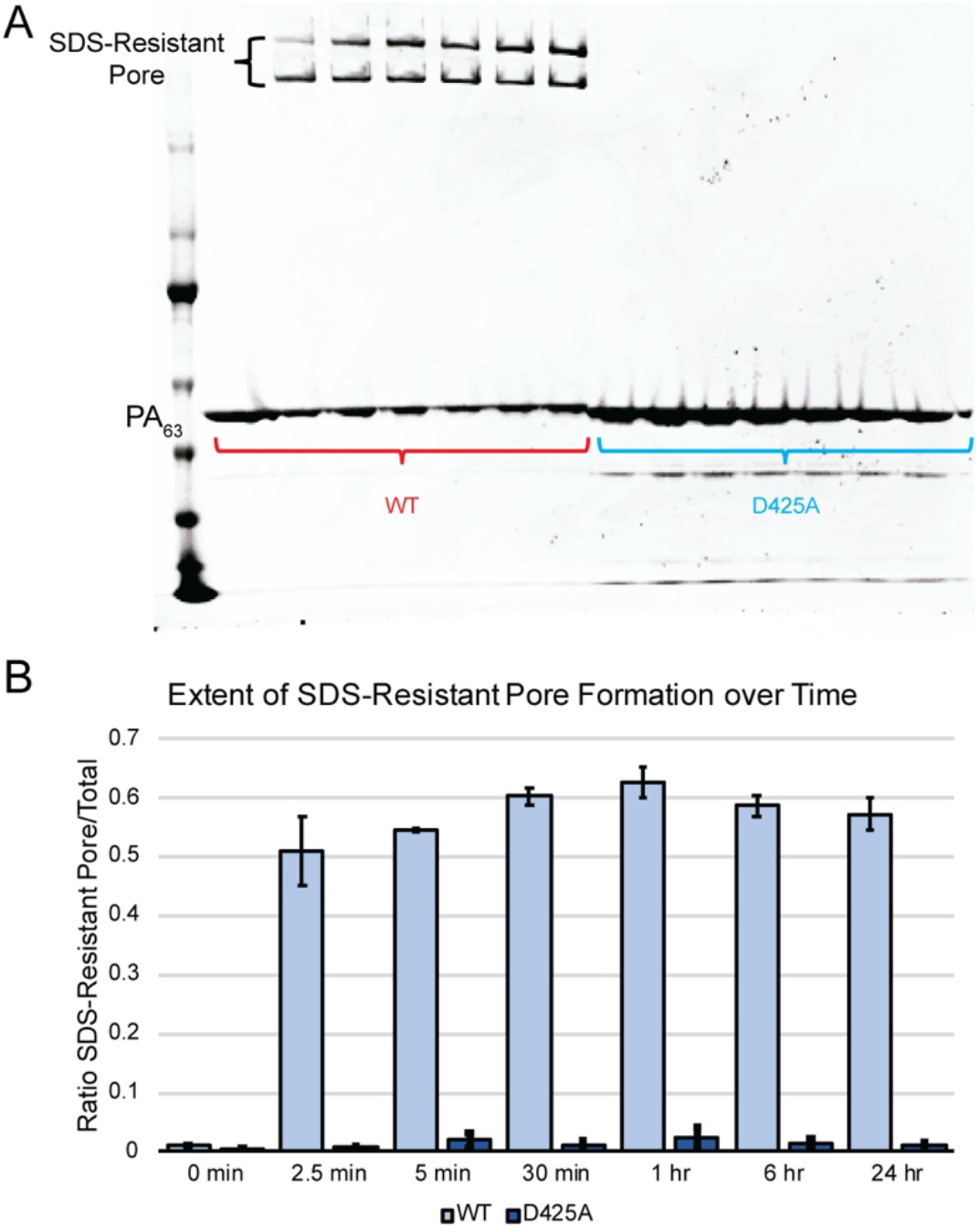
Pore formation assay by SDS-PAGE gel. (**A**) SDS-PAGE analysis of the WT and D425A PA mutant heptamers after exposure to pH 5.5 over a period of 24 hours. (**B**) Quantification of results presented in panel A.

**Figure3-figure supplement 2.**
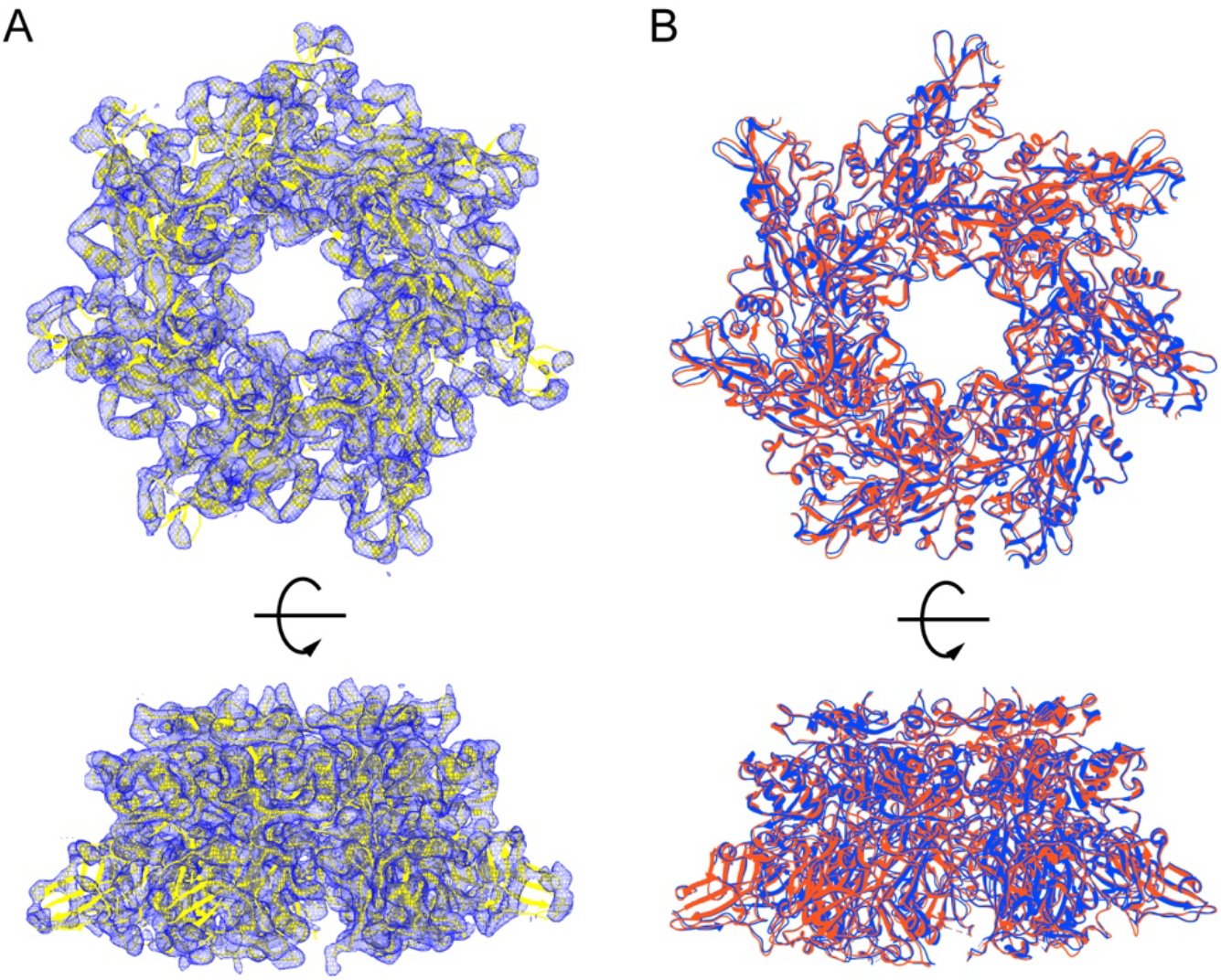
D425A mutant forms heptameric prepore similar to the wild-type at pH 8.5. (**A**) Low-resolution (5.5 Å) X-ray crystal structure of D425A mutant. (**B**) Superposition of the wild-type heptamer structure (red ribbon; PDB entry: 1TZO (Lacy et al., 2004)) with the structure of D425A mutant (blue ribbon) based on the Cα atoms.

**Figure3-figure supplement 3.**
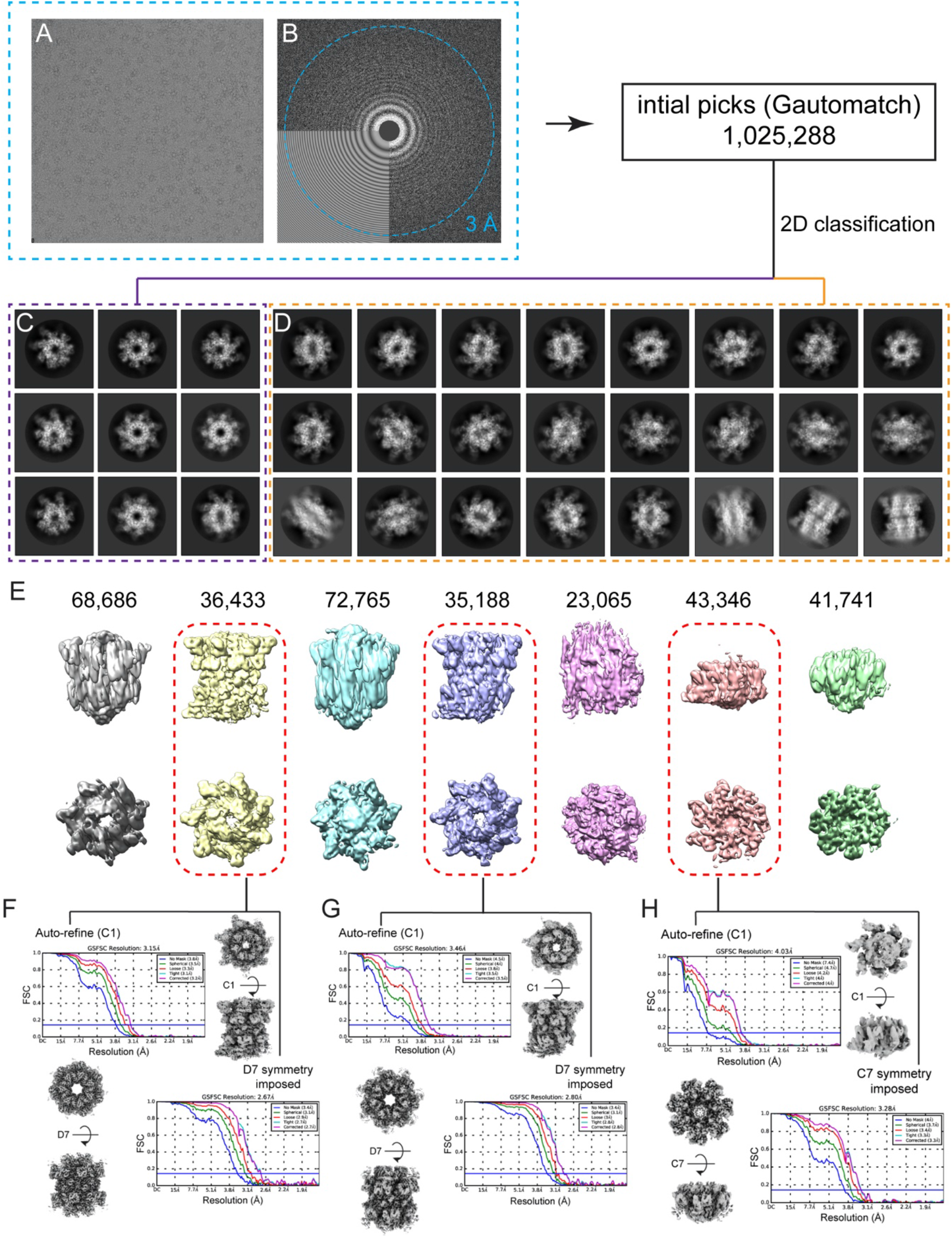
Data collection and image processing workflow for PA D425A mutant at pH 5.5. (**A**) Representative micrograph after alignment with Motioncor2. (**B**) Power spectrum calculated with Gctf, with resolution fit beyond 3.0Å. (**C**) Representative 2D classes of heptamer monomer. (**D**) Representative 2D classes of heptamer dimer. (**E**) 3D classification of D425A at pH 5.5. (**F,G,H**) Refinement of three well resolved cryo-EM maps from the 3D classification. Both C1 and symmetric (C7 or D7) refinement were applied to each class as indicated.

**Figure3-figure supplement 4.**
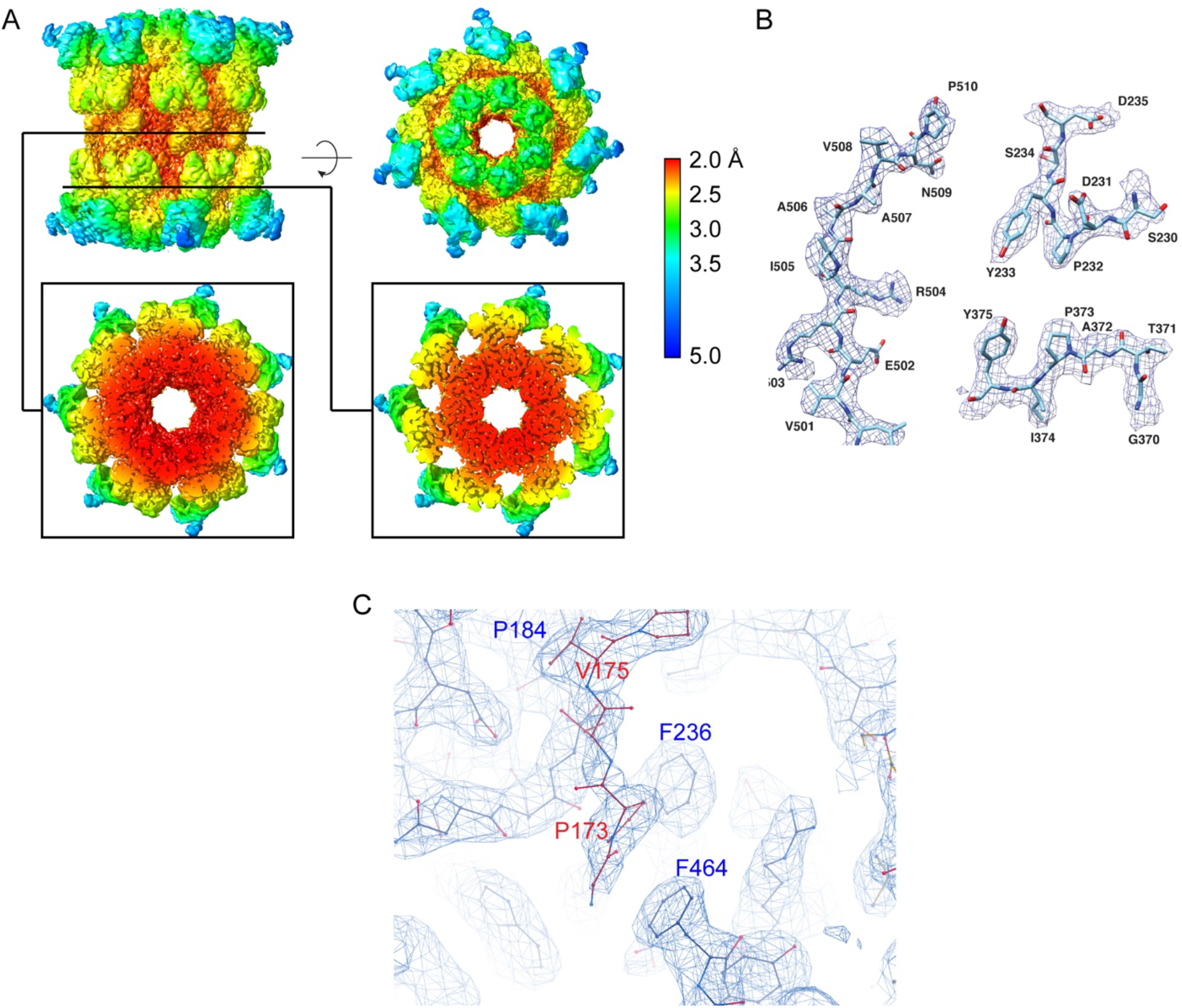
The cryo-EM structure of the D425A pH 5.5 from heptamer dimer. (**A**) Local resolution plot of the reconstruction with D7 symmetry implied. (**B**) Representative regions of structure fit into the density. (**C**) N-terminal sequence of PA loaded to the hydrophobicα-clamp of another PA in the dimeric heptamer.

**Figure3-figure supplement 5.**
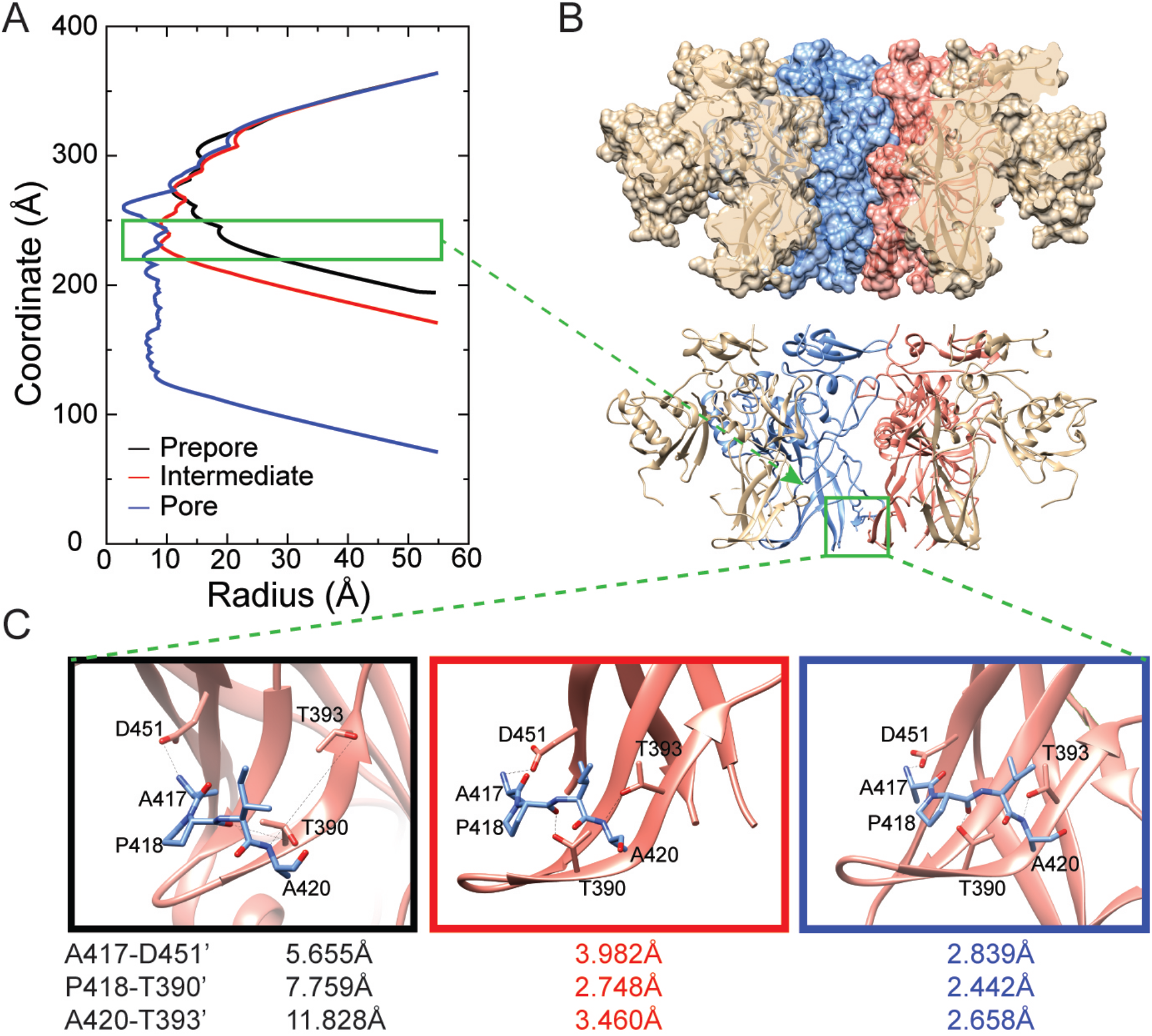
The D425A heptamer structure solved at pH5.5 represents a structural intermediate of the pore maturation pathway. (**A**) Plot of the calculated channel radius through the center of the formed and modeled channels of the WT prepore (black), D425A pH5.5 intermediate (green) and the mature WT pore (blue) states. (**B**) Location of the interaction described in panel C as viewed from the pore lumen. (**C**) Close-up view of the inter-subunit interactions that form during maturation of D425A and WT complexes at low pH. The A_417_-A_420_ loop interaction with the triple-stranded β-sheet (2β_448-458_, 2β_375-385_ and 2β_389-395_) of the adjacent subunit in the WT prepore (left), the D425A intermediate (middle), and the WT pore (right).

**Figure3-figure supplement 6.**
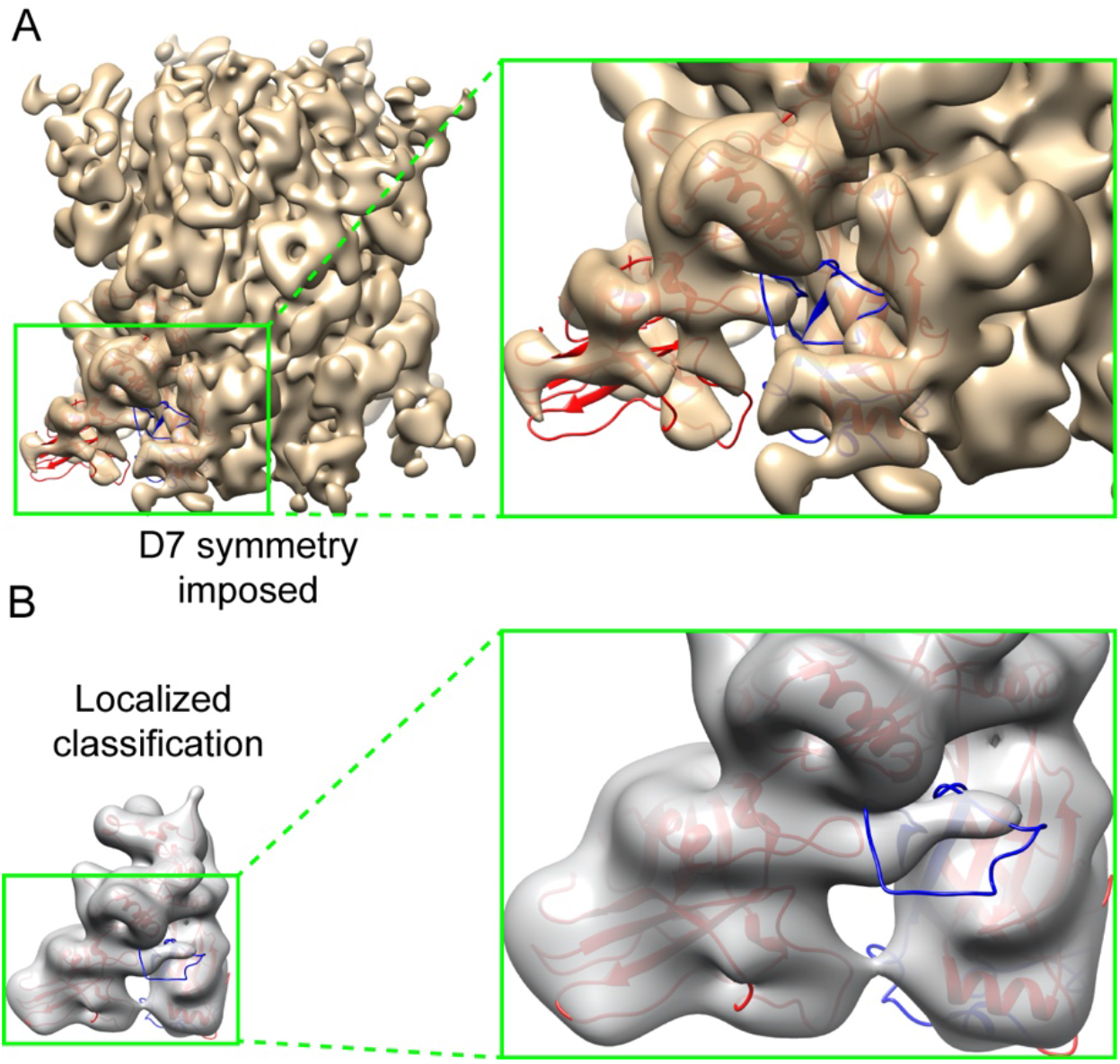
The PFM and domain 4 are resolved in the monomeric D425A pH 5.5 structure with low-pass filtering. (**A**) The D7 symmetrical reconstruction lowpass filtered to 12 Å with a close-up of the PFM and domain 4 of one monomer shows a lack of density to accommodate the PFM (colored blue) displayed in the cartoon diagram. (**B**) Class 1 of the localized reconstruction lowpass filtered to 12 Å with a close-up of the PFM and domain 4 of a monomer identifies density that accommodates the PFM loop, colored blue, and all of domain 4, colored red in bottom left of model, in the cartoon diagram.

**Figure4-figure supplement 7.**
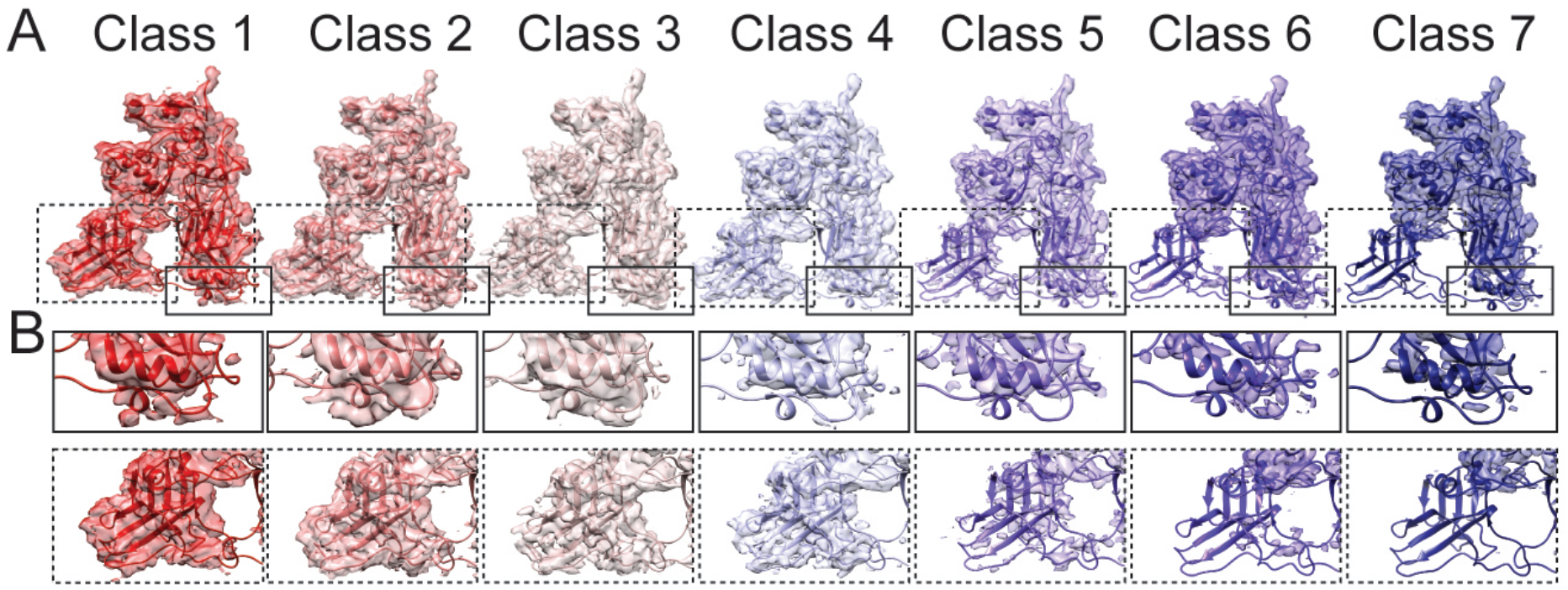
Localized reconstruction of individual monomers within the D425A complex. (**A**) Seven monomeric classes arranged in order of decreasing domain 4 density. (**B**) Close up of the region near the 2α_343-350_ helix (solid boxes) and domain 4 (dashed boxes).

**Supplemental Table 1.** Crystallographic data for D425A at pH 8.5

| PA-63 (D425A) |  |
| --- | --- |
| **Data Collection** |  |
| Unit-cell parameters (Å, °) | $a=166.18$, $b=144.25$, $c=304.83$,<br>$\beta=102.4$ |
| Space group | $P2_1$ |
| Resolution (Å)<sup>1</sup> | 49.78-5.50 (5.69-5.50) |
| Wavelength (Å) | 1.0000 |
| Temperature (K) | 100 |
| Observed reflections | 155,920 |
| Unique reflections | 45,836 |
| $\langle I/\sigma(I) \rangle$<sup>1</sup> | 9.0 (2.2) |
| Completeness (%)<sup>1</sup> | 99.2 (99.8) |
| Multiplicity<sup>1</sup> | 3.4 (3.6) |
| $R_{\text{merge}}$ (%)<sup>1,2</sup> | 9.1 (61.2) |
| $R_{\text{meas}}$ (%)<sup>1,4</sup> | 10.8 (72.2) |
| $R_{\text{pim}}$ (%)<sup>1,4</sup> | 5.8 (38.1) |
| $CC_{1/2}$<sup>1,5</sup> | 0.997 (0.772) |
| **Refinement** |  |
| Resolution (Å)<sup>1</sup> | 44.83-5.50 |
| Reflections (working/test)<sup>1</sup> | 43,559/2,250 |
| $R_{\text{factor}} / R_{\text{free}}$ (%)<sup>1,3</sup> | 25.1/27.8 |
| No. of atoms | 53,490 |
| **Model Quality** |  |
| R.m.s deviations |  |
| Bond lengths (Å) | 0.007 |
| Bond angles (°) | 0.980 |
| $B$-factor (Å<sup>2</sup>) Wilson Plot | 257.0 |
| Coordinate Error (Luzzati Plot, Å) | 0.90 |
| Ramachandran Plot |  |
| Most favored (%) | 89.2 |
| Additionally allowed (%) | 9.3 |
| Outliers (%) | 1.5 |

<sup>1</sup>Values in parenthesis are for the highest resolution shell.
<sup>2</sup> $R_{\text{merge}} = \sum_{hkl} \sum_i |I_i(hkl) - \langle I(hkl) \rangle| / \sum_{hkl} \sum_i I_i(hkl)$ , where $I_i(hkl)$ is the intensity.
<sup>3</sup> $R_{\text{factor}} = \sum_{hkl} ||F_{\text{obs}}(hkl)| - |F_{\text{calc}}(hkl)|| / \sum_{hkl} |F_{\text{obs}}(hkl)|$ ; $R_{\text{free}}$ is calculated in an identical manner using 5% of randomly selected reflections that were not included in the refinement.
<sup>4</sup> $R_{\text{meas}}$ = redundancy-independent (multiplicity-weighted) $R_{\text{merge}}$ (Evans, 2006, 2011). $R_{\text{pim}}$ = precision-indicating (multiplicity-weighted) $R_{\text{merge}}$ (Diederichs and Karplus, 1997; Weiss, 2001).
<sup>5</sup> $CC_{1/2}$ is the correlation coefficient of the mean intensities between two random half-sets of data (Evans, 2012; Karplus and Diederichs, 2012).

**Supplemental Table 2.**
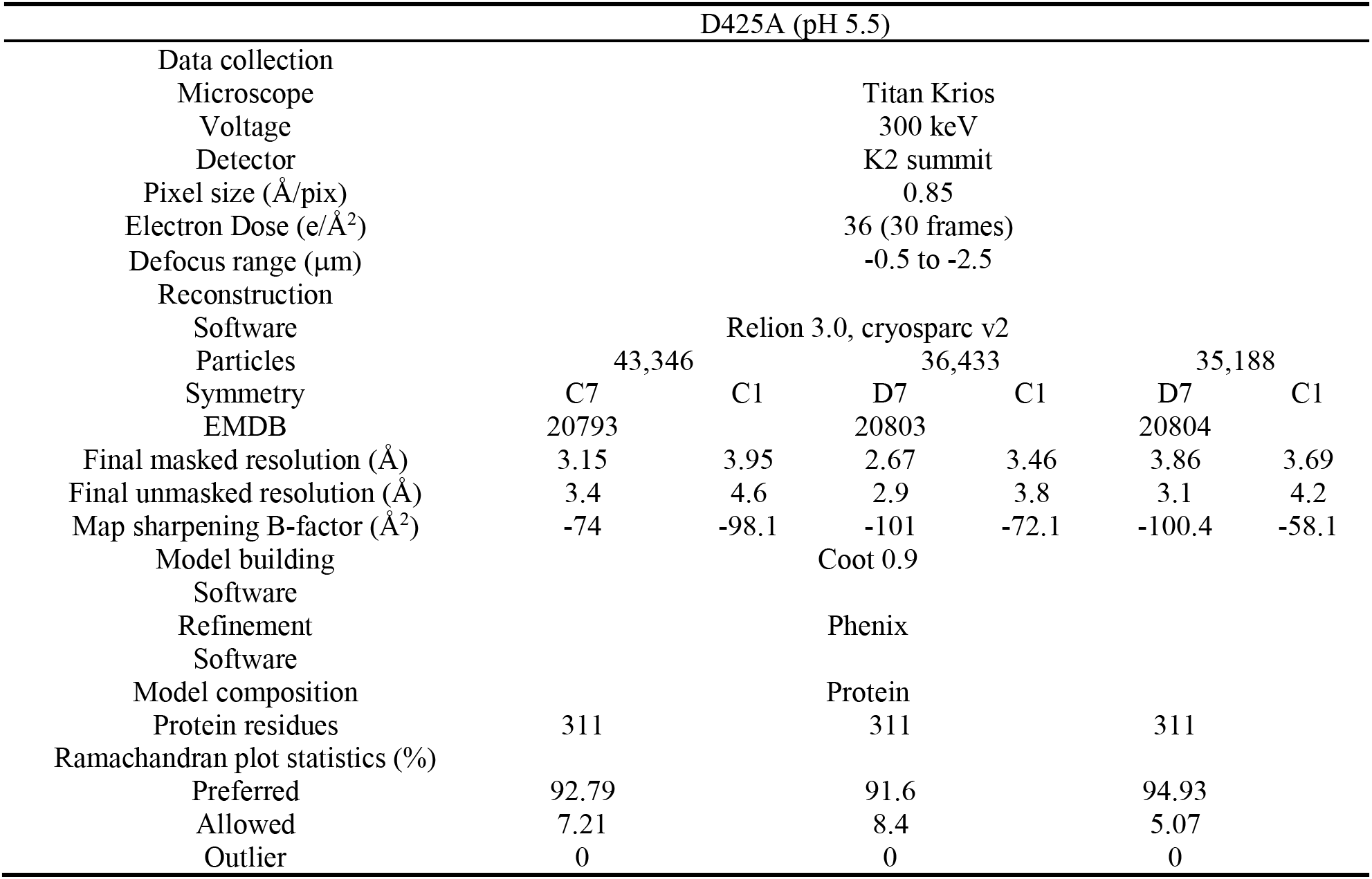
Statistics and model refinement for the D425A Anthrax pore-intermediate solved by cryo-EM

